# High performance anion exchange chromatography purification of probiotic bacterial extracellular vesicles enhances purity and anti-inflammatory efficacy

**DOI:** 10.1101/2023.05.01.538917

**Authors:** Nicholas H. Piroli, Laura Samantha C. Reus, Zuzanna Mamczarz, Sulayman Khan, William E. Bentley, Steven M. Jay

## Abstract

Bacterial extracellular vesicles (BEVs), including outer membrane vesicles (OMVs), have emerged as a promising new class of vaccines and therapeutics to treat cancer and inflammatory diseases, among other applications. However, clinical translation of BEVs is hindered by a current lack of scalable and efficient purification methods. Here, we address downstream BEV biomanufacturing limitations by developing a method for orthogonal size- and charge-based BEV enrichment using tangential flow filtration (TFF) in tandem with high performance anion exchange chromatography (HPAEC). The data show that size-based separation co-isolated protein contaminants, whereas size-based TFF with charged-based HPAEC dramatically improved purity of BEVs produced by probiotic Gram-negative *Escherichia coli* and Gram-positive lactic acid bacteria (LAB). *E. coli* BEV purity was quantified using established biochemical markers while improved LAB BEV purity was assessed via observed potentiation of anti-inflammatory bioactivity. Overall, this work establishes orthogonal TFF + HPAEC as a scalable and efficient method for BEV purification that holds promise for future large-scale biomanufacturing of therapeutic BEV products.

## 1. Introduction

Bacterial extracellular vesicles (BEVs), including the Gram-negative-derived subset of outer membrane vesicles (OMVs), have been applied as vaccines, antimicrobial agents, and therapeutics for a wide variety of immune-associated diseases, such as cancer and inflammatory bowel disease ^1,2^. BEVs are generally defined as being 20-200 nm in diameter and encapsulate bioactive protein and RNA cargo within a protective proteolipid membrane ^3^. While this heterogenous cargo and nanoscale size offers promise for multipotent therapeutic mechanisms of action, these same characteristics present challenges for BEV purification. Commonly-used BEV separation methods leverage biophysical properties such as size, as this distinguishes BEVs from their bacterial producer cells. However, BEV size overlaps with bacterial flagella, pili, and protein aggregates, and thus separation approaches involving ultracentrifugation (UC) can co-isolate many impurities. This can be addressed somewhat by employing a density gradient ^4^, but this has limited scalability. Alternatives such as tangential flow filtration (TFF) and size exclusion chromatography (SEC) are more scalable than UC, but the potential for co-isolation of impurities remains. Importantly, these impurities are potentially immunogenic ^5-7^, even in trace quantities, and thus present safety and efficacy concerns, especially for anti-inflammatory probiotic BEV therapeutics ^8^. Thus, overall there is a significant need for improving downstream BEV biomanufacturing.

Anion exchange chromatography (AEC) separates molecules based on differences in net charge and is widely used in industrial biomanufacturing within automated high performance liquid chromatography systems (HPLC). AEC relies on a positively-charged matrix to bind negatively-charged molecules while neutral or positively-charged molecules are washed away, followed by elution of bound molecules with a high ionic strength buffer. AEC has recently been applied for mammalian EV purification, enabled by the intrinsic negative membrane surface charge of EVs ^9,10^. Like EVs, BEVs from virtually all bacteria possess a negative membrane surface charge ^11^, and thus we hypothesized that AEC is a potential solution for scalable, non-size-based BEV purification.

Here, we report that high performance AEC (HPAEC) in tandem with size-based TFF dramatically improves probiotic *Escherichia coli* Nissle 1917 (EcN) BEV purity compared to size-based methods alone as indicated by reduced flagellin protein impurities. Since flagellin is negatively charged at physiologic pH, we optimized HPAEC by lowering pH to preferentially purify BEVs from negatively-charged free protein. Finally, our method for orthogonal TFF and HPAEC was successfully applied to BEVs produced from several well-characterized Gram-positive probiotic bacteria (*Lacticaseibacillus rhamnosus, Lacticaseibacillus casei*, and *Limosilactobacillus reuteri)*, as indicated by retained (*L. reuteri)* or potentiated (*L. casei, L. rhamnosus*) anti-inflammatory bioactivity. Given the widespread use of HPAEC in industry, we anticipate that this approach could be readily adapted for scalable biomanufacturing of probiotic BEVs, potentially enabling the rapid expansion of an emerging class of biotechnology products that have utility as vaccines and therapeutics against infections, chronic inflammatory diseases, and cancer ^12-17^.

## 2. Materials and Methods

### 2.1 Bacterial culture and production

The probiotic *Escherichia coli* Nissle 1917 (EcN) was obtained from Mutaflor (Canada). EcN was transformed with an arabinose-inducible expression vector encoding a ClyA-sfGFP fusion protein (pBAD-ClyA-sfGFP) to achieve GFP loading into BEVs, as previously described ^18^. EcN was cultured at 37□°C in 200 mL Luria-Bertani broth (LB) supplemented with 0.2% arabinose for 16 hours with 250 rpm shaking before BEV harvesting. All lactic acid bacteria (LAB) strains were obtained through ATCC (*Lacticaseibacillus rhamnosus* GG (ATCC 53103), *Lacticaseibacillus casei* (ATCC 393), *Limosilactobacillus reuteri* F 275 (ATCC® 23272™). LAB strains were cultured aerobically in 400 mL MRS broth at 37□°C without shaking for 24 hours before BEV harvesting. All BEV samples were initially processed using centrifugation of culture followed by TFF of sterile culture supernatant, as detailed below. Additionally, after the terminal isolation procedure, all BEV samples were sterile filtered with a 0.2 uM syringe filter and stored at -20 □°C for no longer than 4 weeks until use.

### 2.2 Tangential flow filtration (TFF)

First, sterile culture supernatant was generated by centrifuging culture media at 10,000 x *g* x 10 minutes at 4□°C to remove cells. The supernatant was transferred to a new tube and centrifuged again at 10,000 x *g* x 20 minutes at 4□°C to remove residual cells and debris. Then, supernatant was filtered through a 0.45 um polyethersulfone (PES) bottle top filter prior to TFF; TFF was performed using a using a KrosFlo KR2i TFF system (Spectrum Labs, Los Angeles, CA, USA) equipped with a 300-kDa MWCO hollow fiber filter composed of a modified PES membrane (D02-E100-05-N; Spectrum Labs). Each filter was used no more than ten times, cleaned by flushing with 100 mL dH20 followed by circulating 0.5M NaOH for 30 minutes, and stored in 20% ethanol. Flow rate was set at 106 mL/min to maintain a shear rate of 4000 s^−1^ and backflow pressure was automatically adjusted to maintain a transmembrane pressure of 5 psi.

Sterile supernatant was initially concentrated to 25 mL followed by diafiltration with 5-fold volumes of 1× PBS and then finally concentrated to 7-10 mL. BEV-depleted conditioned media samples were obtained by collecting sub-300 kDa permeate during the initial concentration step after the first 10 mL of permeate was discarded. FinallTFF-isolated BEV samples were concentrated to 1 mL using 100 kDa ultrafiltration columns, sterile filtered with 0.2 uM PES syringe filters, and stored at -20□°C for no longer than 4 weeks before use.

### 2.3 Size-exclusion chromatography (SEC)

TFF-isolated BEV samples were further processed with SEC using qEV Original columns (ICO-35; Izon Science) per the manufacturer’s protocol. For some experiments Sepharose CL-6B beads (CL6B200, Sigma-Aldrich) were packed into a 10 mL column of identical dimensions as qEV columns and SEC was performed with the same procedure. After flushing the SEC columns with 1× PBS, 0.5 mL of the TFF-isolated BEV sample was applied to the top of the column and a 2.0 mL fraction after the void volume was collected. For some experiments in Figure 1, either a greater number of sequential fractions or 0.5 mL fractions were collected. The BEV-containing 2.0 mL fractions were then concentrated using 300 kDa MWCO ultrafiltration columns to a final volume of 0.5 mL.

**Figure 1.**
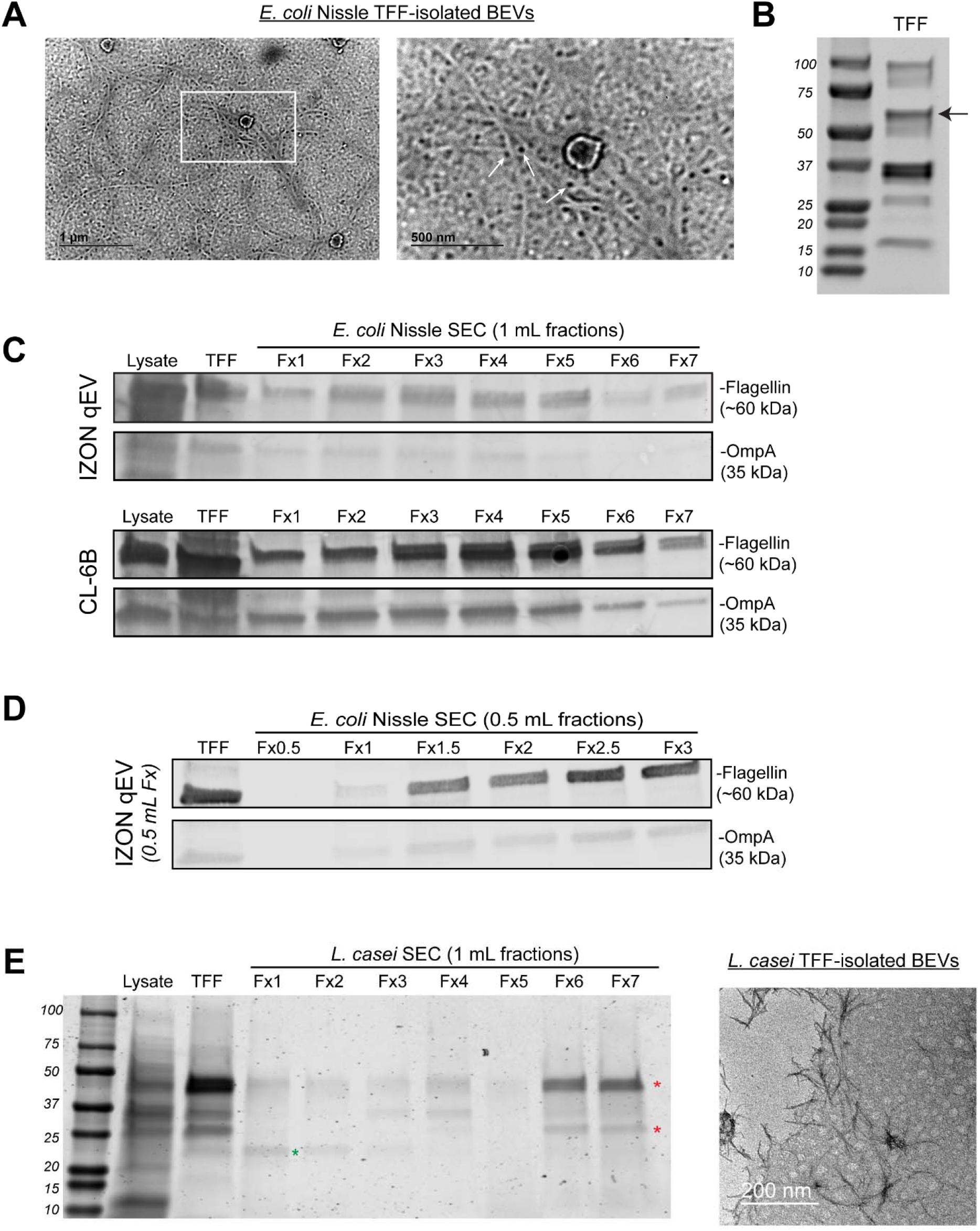
Size-based methods for BEV separation co-isolate flagellin impurities. **A)** anti-flagellin immunogold TEM of EcN BEVs purified by TFF only; scale bars 1 um (left) and 500 nm (right, inset), **B)** Coomassie-stained SDS-PAGE of EcN BEVs purified by TFF only, **C)** Western blot analysis of relative levels of OmpA (BEV marker) and flagellin (co-isolated impurity) in EcN cell lysate, TFF-purified BEVs, and sequential 1mL fractions of additional SEC purification of previously TFF-purified BEVs with either Izon qEV (upper blots) or Sepharose CL-6B (lower blots). **D)** western blot analysis of relative OmpA and flagellin levels following Izon qEV purification with reduced 0.5 mL fraction volume. **E)** Coomassie-stained SDS-PAGE of *L. casei* cell lysate and BEVs purified by TFF only, or TFF + SEC (left) and transmission electron micrographs of TFF-purified *L. casei* BEVs, scale bar 200 nm.

### 2.4 Anion Exchange Chromatography

Benchtop AEC was performed using a weak anion exchanger resin (DEAE Sepharose fast flow; Cytiva, 17070901) packed in a 10 mL column. TFF-purified BEVs were buffer exchanged with 20 mM Tris-HCl prior to loading into AEC columns. AEC columns were first equilibrated with five column volumes (CVs) of 20 mM Tris-HCl (equilibration buffer), then 0.5 mL of TFF-isolated BEVs were added directly onto the AEC resin and allowed to incubate at room temperature for 1 hour to promote maximal binding. Then, the column was washed with three CVs of 20 mM Tris-HCl, followed by elution with five CVs 1 M NaCl in 20 mM Tris-HCl (elution buffer). Fractions were collected and concentrated using 100 kDa MWCO ultrafiltration columns.

High performance AEC was performed using a Bio-Rad NGC Quest 10 Plus Chromatography System (Bio-Rad; 7880003) equipped with a 5 mL HiTrap Q column (Cytiva; 17115401) that contains a strong anion exchange resin to support AEC at reduced pH. The UV-vis spectrophotometer was set to detect wavelengths of 210 nm, 215 nm, 220 nm, and 280 nm. For pH 7.4 experiments, the equilibration buffer was 20 mM Tris-HCl and the elution buffer was 20 mM Tris-HCl with 1 M NaCl. For pH 5.8 experiments, the equilibration buffer was 20 mM piperazine and the elution buffer was 20 mM piperazine with 1 M NaCl. Following column equilibration with equilibration buffer, 1E11 particles of TFF-purified BEVs suspended in 5 mL of appropriate equilibration buffer were loaded into the column at 0.1 mL per minute. Then, the column was washed with seven CVs of equilibration buffer before elution commenced beginning with two CVs of 0.1 M NaCl, then two CVs of 0.2 M NaCl, and onwards up to 1 M NaCl in steps of 0.1M. Fractions corresponding to each NaCl concentration were collected using an automated fraction collector.

### 2.5 BEV characterization

Size distribution and particle concentration were assessed using a NanoSight LM10 (Malvern Instruments) with Nanoparticle Tracking Analysis (NTA) software, version 2.3. For each sample, three 30-second videos were captured with a camera level set at 14. BEV samples were diluted to obtain 20–100 particles per frame and at least 200 completed tracks per video to ensure accurate analysis. The detection threshold was set at 3 and kept constant across all replicates and samples. Total protein concentration in BEV samples was determined using Bicinchoninic acid assay (BCA; 786-571; G-Biosciences). AEC samples were buffer exchanged with 1× PBS prior to BCA to remove excess NaCl and adjust pH to 7.4.

### 2.6 Western blot

EcN BEV purity was assessed by Western blot analysis of relative levels of flagellin and the BEV marker OmpA. 40 uL of BEV samples were loaded into 4–15% Mini-PROTEAN® TGX SDS-PAGE gel, except TFF-only BEV samples which were limited to 30 ug total protein due to high protein concentration. The primary antibodies were rabbit anti-OmpA (1:25,000 dilution; 117097; Antibody Research Corporation) and rabbit anti-flagellin (1:1000 dilution; ab93713; Abcam). A 1:10,000 dilution of a goat anti-rabbit secondary antibody (926-32211; LI-COR Biosciences) was used. Antibodies were diluted in a 1:1 ratio of Odyssey blocking buffer (LI-COR; # 927-40000) and PBS + 0.4% Tween 20. Protein bands were imaged using a LI-COR Odyssey CLX Imager. Densitometry was utilized to determine relative ratios of OmpA:flagellin using the associated software. Comparisons of OmpA:flagellin ratios were only made between samples processed in parallel, derived from the same batches of culture, and run on the same gel and blot.

### 2.7 Transmission electron microscopy (TEM)

Morphology of BEVs was visualized via TEM using negative staining. First, 20 uL of BEV sample was fixed in 2% EM-grade paraformaldehyde (157-4-100; Electron Microscopy Sciences, Hatfield, PA, USA) for 30 min at room temperature. For all subsequent steps, ultra-thin carbon-coated copper grids were floated carbon-film down on droplets of regents placed on a sheet of parafilm with blotting of excess liquid between steps. To adhere fixed BEVs, grids were floated on a droplet of PFA-BEV mixture for 20 minutes. The BEV-adhered grid was then briefly washed with PBS and floated on a droplet of 1% glutaraldehyde (in 1× PBS) for 5 minutes. Next, the grid was washed 5 times with dH20 (2 minutes per wash), and then negative stained using uranyl-acetate replacement stain (22405; Electron Microscopy Sciences) for 10 minutes. The grids were allowed to dry overnight before imaging on a JEOL JEM 2100 LaB6 TEM with a 200 kV accelerating voltage.

For immunogold TEM, the BEV-coated grids were prepared as above except immunostaining was performed prior to negative staining with uranyl acetate replacement stain. For immunostaining, free aldehyde groups in BEV-coated grids were quenched with 50 mM glycine dissolved in PBS for 15 minutes. Then, grids were washed 3x with dH20 and blocked with Odyssey Blocking Buffer (927-40000; LI-COR) for 30 minutes. Following 3x wash with PBS, grids were floated on rabbit anti-flagellin polyclonal antibody (ab93713; Abcam) diluted 1:100 in blocking buffer + 0.1% Tween 20 overnight at 4□°C. Then, grids were washed 3x with blocking buffer + 0.2% Tween and floated on biotin-conjugated anti-rabbit secondary antibody (ab6720; Abcam) diluted 1:100 in blocking buffer + 0.1% Tween 20. Following 3x PBS wash, grids were floated on streptavidin conjugated 5 nm gold nanoparticles for 30 minutes, then washed 3x with PBS, and fixed with 1% glutaraldehyde for 15 minutes. Finally, grids were washed 3x with PBS, negative stained with uranyl-acetate replacement stain for 10 minutes, then blotted and allowed to dry overnight before imaging.

### 2.8 Zeta potential

For Zeta-Potential (ζ-potential) measurement, phase analysis light scattering (PALS) measurements were performed using a NanoBrook Omni particle analyzer (Brookhaven Instruments Corporation). BEVs were isolated with TFF and concentrated to approximately 1E11 particles/mL in PBS pH 7.4, then diluted 10-fold with dH20 (final concentration 0.1× PBS). Diluted BEVs were loaded into disposable cuvettes, and three technical replicates were collected at 25□°C using Henry equation analysis.

### 2.9 Macrophage inflammatory assay

To compare and verify probiotic BEV anti-inflammatory bioactivity between isolation methods, we utilized a mouse macrophage stimulation assay ^19^. RAW264.7 mouse macrophages were seeded at 75,000 cells/well in a 48-well plate in DMEM + 5% FBS + 1% penicillin/streptomycin (Media). Then, 24 h later, cells were pretreated with media supplemented with either: i) 1× PBS (6 wells total), ii) BEV-depleted conditioned media (permeate from TFF), iii) 1 μg/mL dexamethasone as a positive control (Dex; D4902-25 MG; Sigma-Aldrich), and iv) BEV groups from various isolation methods (TFF only, TFF + SEC, or TFF + HPAEC). All groups were performed in triplicate and doses between BEV-depleted conditioned media and BEV groups were normalized by protein content. After 24 h of incubation, the pre-treatments were removed and 10 ng/mL LPS (resuspended in 1× PBS; L4391-1MG; Sigma-Aldrich) was added to all groups except three PBS pretreated wells (media only group) to stimulate inflammatory responses. Conditioned media was collected 4 hours post-LPS stimulation, stored at -80°C, and levels of tumor necrosis factor-α (TNF-α) were assessed by ELISA within 3 days (DY410; R&D Systems). The cytokines TNF-α, IL-6, and IL-10 were previously screened and TNF-α was selected due to a high dynamic range of dose-dependent suppression following BEV pretreatment.

## 3. Results

### 3.1 Size-based methods for BEV separation co-isolate flagellin impurities

TFF and SEC are scalable alternatives to ultracentrifugation for separation of both mammalian and bacterial EVs that should enable increased purity. In Figure 1, however, we observed flagella and its constituent protein subunit flagellin were co-isolated when applying TFF purification of *E. coli* Nissle 1917 (EcN) BEVs. Immunogold electron micrographs of TFF-purified BEVs revealed abundant threadlike fibers marked by 5 nm gold nanoparticles conjugated to anti-flagellin antibody (Figure 1A). Additionally, Coomassie-stained SDS-PAGE revealed a prominent ∼60 kDa band that corresponds to the expected molecular weight of flagellin (Figure 1B, black arrow). Thus, TFF co-isolates relatively large quantities of flagellin, possibly due to BEVs and flagella or aggregates of flagellin sharing similar size ranges. Next, we investigated if SEC purification of the TFF-purified BEVs could reduce co-isolated flagellin. Western blot analysis of the BEV marker OmpA and flagellin revealed only marginal reductions in relative amounts of flagellin compared to OmpA with Izon qEV SEC purification. Similar results were observed for an alternative SEC resin, Sepharose CL-6B (Figure 1C), and reducing fraction sizes from 2 mL to 0.5 mL also did not result in improvement in BEV:flagellin ratio (Figure 1D). Thus, size-based purification alone, either with TFF only or TFF + SEC, has a limited capacity to reduce certain co-isolated proteins, particularly flagellin.

To assess if the observed results might be specific to EcN, we next tested TFF and TFF + SEC purification of BEVs from the Gram-positive probiotic *L. casei*. Since *L. casei* BEVs lack reliable markers with commercially-available antibodies^11^, we used Coomasie-stained SDS-PAGE and electron microscopy to assess purity. Like EcN BEVs, there were several major protein bands in the TFF sample (Figure 1E). Further purification with SEC revealed several protein bands eluted in non-BEV containing fractions, suggesting their identity as impurities co-isolated with TFF (Figure 1E indicated by red stars). In contrast, a ∼30 kDa band eluted only in BEV-containing fractions (green star) suggesting its identity as a *L. casei* BEV marker. Notably, a significant fraction of the co-isolated proteins remained in the BEV-containing SEC fraction, similar to results observed with flagellin in EcN BEV purification. Additionally, electron micrographs of TFF + SEC-purified *L. casei* BEVs revealed the presence of dark staining fibrous aggregates similar in size to BEVs that may represent protein aggregates (Figure 1E). This is likely evidence that size-based purification co-isolated proteins in *L. casei* BEV samples, similar to EcN.

Overall, these results demonstrated that size-based purification with TFF or TFF + SEC coisolate protein impurities. For EcN, we identified flagellin as one such co-isolated impurity present in relatively high concentrations in both TFF and TFF + SEC processed BEV samples. Furthermore, we identified potential co-isolated impurities in *L. casei* BEV samples. Since coisolated protein was consistently detected across multiple size-based purification schemes, we decided to explore non-size-based purification methods.

### 3.2 Anion exchange chromatography (AEC) can be used for BEV separation

As noted above, AEC purifies molecules by charge and therefore we hypothesized that it could be used in tandem with size-based methods yielding an orthogonal methodology for BEV separation. First, we confirmed prior reports that BEVs from EcN, *L. reuteri, L. casei* and *L. rhamnosus* have a negative surface charge at pH 7.4 (as indicated by negative zeta potential), suggesting the potential for charge-based purification by AEC (Figure 2A,B). Indeed, we found that TFF-isolated EcN BEVs could be recovered from AEC resin following elution with high salt buffer (Figure 2C). The mode particle sizes for TFF-only and TFF + AEC EcN BEVs were similar (148 nm ± 2.3 vs. 140.7 nm ± 7.4). Interestingly, the mean size of AEC-purified EcN BEVs was far lower than those purified by TFF-only (163.5 nm ± 6.5 vs. 229.6 nm ± 7.8). This was due to reduced numbers of very large >300 nm presumably non-BEV particles, whose origins are unknown at present. Regardless, nanoparticle tracking analysis revealed high concentrations of EcN BEVs in AEC elution fractions and low concentrations in wash fractions, suggesting high EcN BEV binding efficiency to AEC resin (Figure 2C).

**Figure 2.**
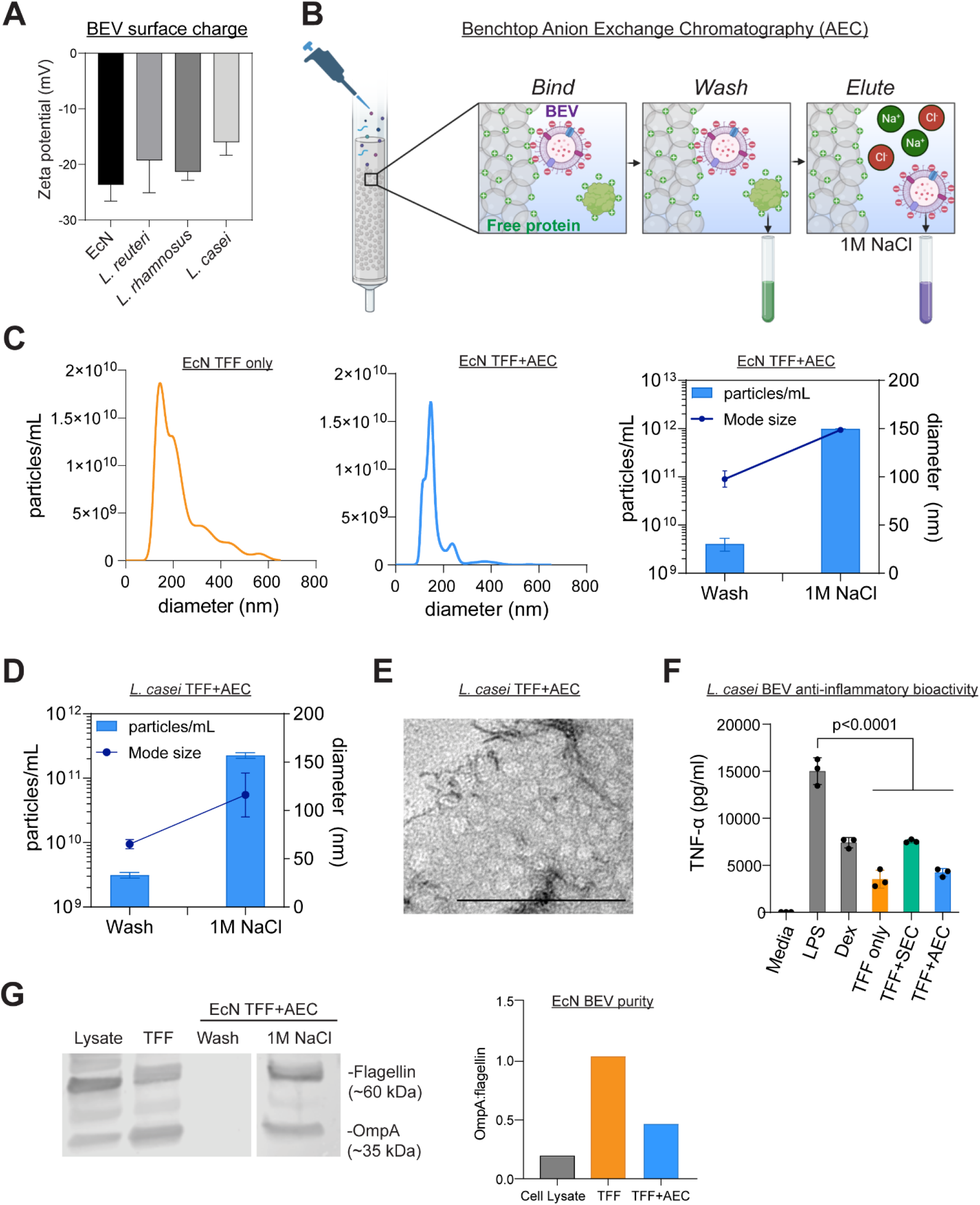
Anion exchange chromatography (AEC) can be used for BEV separation. **A)** Zeta potential of TFF-isolated BEVs from EcN, *L. reuteri, L. rhamnosus* and *L. casei* obtained by phase analysis light scattering (n=3 technical replicates), **B)** schematic depicting benchtop AEC BEV purification process. **C-D)** Nanoparticle tracking analysis of EcN (C) and *L. casei* (D) BEV samples following TFF-only purification or TFF + AEC purification (n=3 technical replicates). **E)** Transmission electron micrographs of *L. casei* BEVs purified by TFF and AEC, **F)** TNF-α levels from RAW264.7 mouse macrophage conditioned media following treatment with dexamethasone (Dex) or equal protein doses of BEVs purified by TFF-only, TFF + SEC, or TFF + HPAEC and subsequent LPS-stimulation (n=3). Data analyzed by one-way ANOVA with Holm-Šidák post hoc test. **G)** EcN BEV purity analyzed using relative levels of OmpA and flagellin to produce an OmpA:flagellin ratio (right) indicative of relative BEV purity of TFF-only and TFF + HPAEC purification.

Next, we tested if AEC could be applied to LAB. Since BEVs from LAB lack protein markers like the flagellin of EcN, we employed an anti-inflammatory bioactivity assay to track AEC purification of *L. casei* BEVs. In *L. casei* AEC elution fractions, we observed high particle counts (Figure 2D), presence of BEVs in electron micrographs (Figure 2E), and anti-inflammatory bioactivity (Figure 2F), altogether indicating that AEC can separate *L. casei* BEVs. Inspired by these results, we sought to characterize EcN BEV purity using ratios of OmpA to flagellin. OmpA is an abundant outer membrane protein that is not naturally secreted except within BEVs and therefore is widely used as an *E. coli* BEV marker, whereas flagellin is an extracellular protein that is not a natural component of BEVs but is commonly co-isolated ^20^. Thus, higher ratios of OmpA:flagellin would indicate greater BEV purity. However, we observed a reduced OmpA:flagellin ratio with AEC purification (Figure 2G). Additionally, dark staining fibrous aggregates were visualized in *L. casei* electron micrographs (Figure 2E), suggesting that non-BEV-associated proteins were also binding to the AEC resin, most likely due to their net negative charge.

### 3.3 Optimized AEC in tandem with TFF improves BEV purification

The results in Figure 2 follow from the fact that flagellin and other proteins are negatively charged at physiologic pH and thus would be eluted with BEVs in the bulk collection scheme that we employed. However, there is a difference in net charge between these entities that we hypothesized could be exploited for improved BEV separation. To test this hypothesis, we first attempted to elute BEVs and flagellin at different times by precisely controlling the ionic strength of elution buffer. This was accomplished using a HPLC system to conduct AEC (HPAEC). Additionally, we utilized an engineered EcN strain that expresses a ClyA-sfGFP fusion protein to direct GFP loading into BEVs to enable fluorescent tracking of BEVs in elution fractions ^15^. We loaded TFF-purified EcN-ClyA-sfGFP BEVs into HPAEC and eluted BEVs with a stepwise [NaCl] gradient beginning at 0.1 M NaCl and increasing to 1 M in 0.1M steps. Fraction analysis revealed particle counts gradually decreased with increasing [NaCl], whereas mode particle size was relatively constant (Figure 3A). Additionally, fluorescence generally tracked with particle counts, with gradually decreasing fluorescence with increasing [NaCl] (Figure 3A). Finally, electron micrographs revealed the presence of BEVs with expected cup-shaped morphology (Figure 3B).

**Figure 3.**
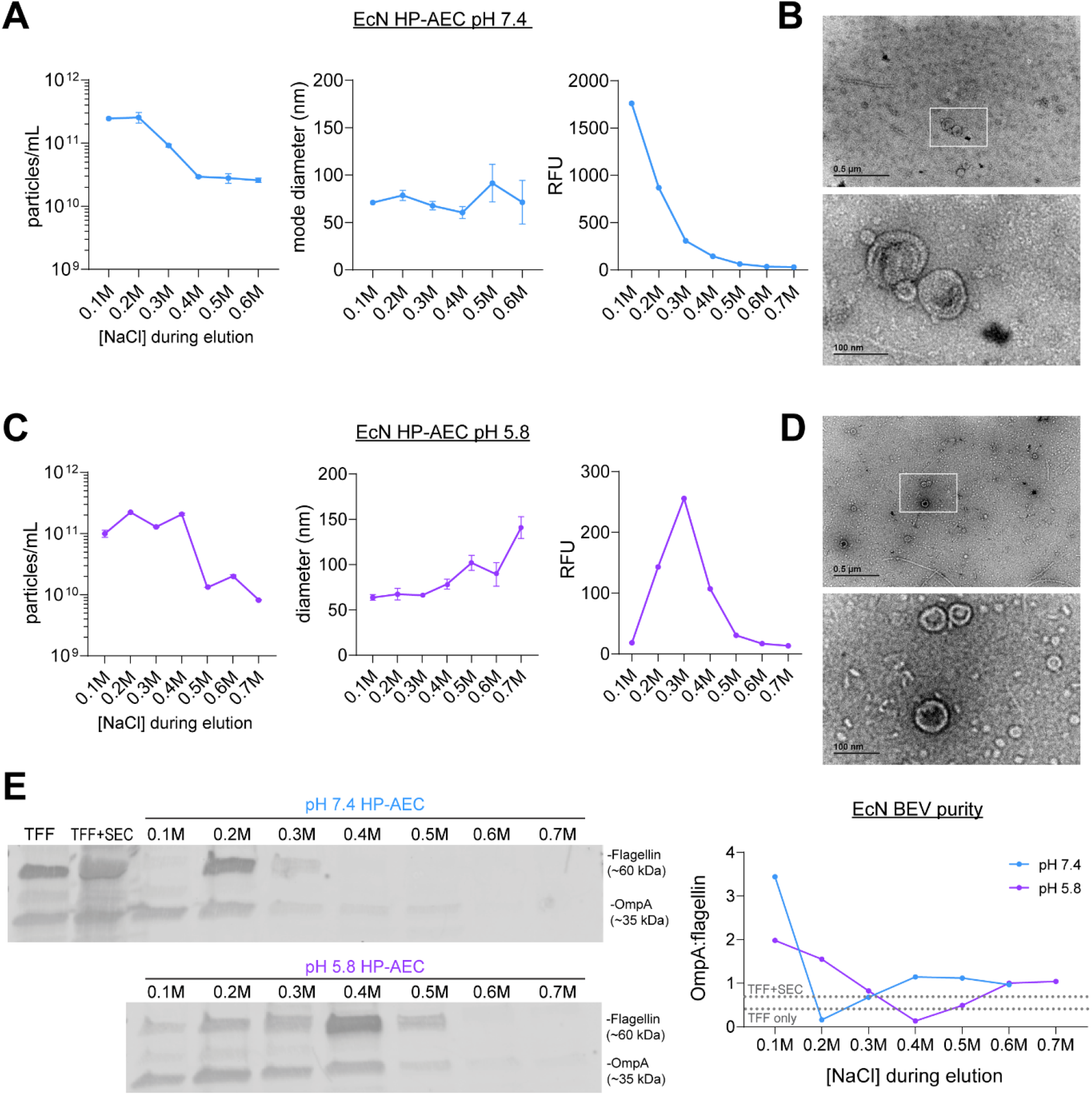
Optimized AEC in tandem with TFF improves BEV purification. **A)** Nanoparticle tracking analysis (left) and fluorescence readings (right) of GFP-loaded BEVs following TFF + HPAEC purification at pH 7.4 (n=3 technical replicates), **B)** transmission electron micrographs of TFF + HPAEC purified BEVs, scale bar 0.5 um (upper) and 100 nm (lower, inset). **C-D)** same analysis as in A-B above except HPAEC was performed at pH 5.8. **E)** Western blot analysis of relative levels of OmpA and flagellin in elution fractions following HPAEC purification at pH 7.4 or pH 5.8, with densitometry quantification of OmpA:flagellin ratio (right).

Additionally, we pursued a strategy involving active modification of the net charge of flagellin via pH adjustment. The negative charge of BEVs likely arises primarily from phosphate groups (within LPS and phospholipids) ^21^, whereas the negative charge of bacterial proteins arises solely from amino acid carboxylic acid side groups. Since the pKa of phosphate groups is less than amino acid carboxylic acid groups, we hypothesized that pH reduction would preferentially preserve the negative charge of BEVs while causing free proteins to lose their negative charge due to protonation. Thus, BEVs would still bind to the AEC resin, whereas protonated free proteins would not bind, or bind more weakly. We tested this hypothesis using BEVs suspended in buffer with pH of 5.8. After HPAEC separation, we observed consistently high particle counts in the 0.1 M through 0.4 M NaCl fractions that decreased 10-fold in later fractions (Figure 3C). Fluorescence readings were also generally high in the 0.1 M through 0.4 M fractions (Figure 3C), but it is worth noting that the low pH likely quenched some sfGFP fluorescence ^22,23^.

Electron micrographs confirmed the presence and expected cup-shaped morphology of BEVs in pH 5.8 AEC samples (Figure 3D). Finally, we quantitatively compared BEV purity between purification methods and pH. Consistent with earlier experiments, the OmpA:flagellin ratio for TFF-only was low (0.3) and increased only slightly with additional SEC purification (OmpA:flagellin = 0.5) (Figure 3E). In contrast, we observed dramatic increases in BEV purity in several of the HPAEC fractions. The greatest overall purity was found in the pH 7.4 0.1 M NaCl fraction (OmpA:flagellin = 3.5) (Figure 3E). However, all other pH 7.4 fractions were characterized by relatively high flagellin (0.2 M and 0.3 M NaCl) or low BEV yield (0.4 M onwards) (Figure 3E). In contrast, pH 5.8 produced several fractions with high purity (0.1 M through 0.3 M) that in total contained 2-fold more particles than the high purity pH 7.4 0.1M fraction (Figure 3E).

### 3.4 HPAEC can be applied for separation of BEVs from Gram-positive lactic acid probiotic bacteria

From the data in Figure 3, we conclude that i) HPAEC in tandem with TFF is superior to size-based methods alone in eliminating co-isolated flagellin impurities in EcN, and ii) pH adjustment in HPAEC is a potential approach for separation of EcN BEVs from negatively charged protein impurities. To test whether HPAEC could be applied to enrichment of BEVs beyond EcN, we used Gram positive lactic acid bacteria probiotics *L. casei, L. reuteri*, and *L. rhamnosus* as BEV producers, all of which have well-characterized BEV efficacy in a variety of diseases ^24-27^. Purification of BEVs from Gram-positive bacteria is complicated in that there are 10-100-fold lower BEV yields and greater numbers of proteins secreted when compared to Gram-negative bacteria ^11^. We found that HPAEC successfully purified BEVs from all three bacteria, as indicated by particle count and electron micrographs (Figure 4A-D). As expected, we observed relatively low UV-vis absorbance during elution and 10-100-fold lower particle counts compared to EcN, although *L. rhamnosus* yields were 10-fold greater than *L. casei* or *L. reuteri* (Figure 4A-C). This could potentially be explained by the slightly more negative zeta potential of *L. rhamnosus* compared to *L. casei* and *L. reuteri*. Additionally, like EcN, particle counts generally declined with increasing [NaCl]. Unlike EcN, however, we observed gradually increasing particle diameter with increasing [NaCl] (Figure 4A-C).

**Figure 4.**
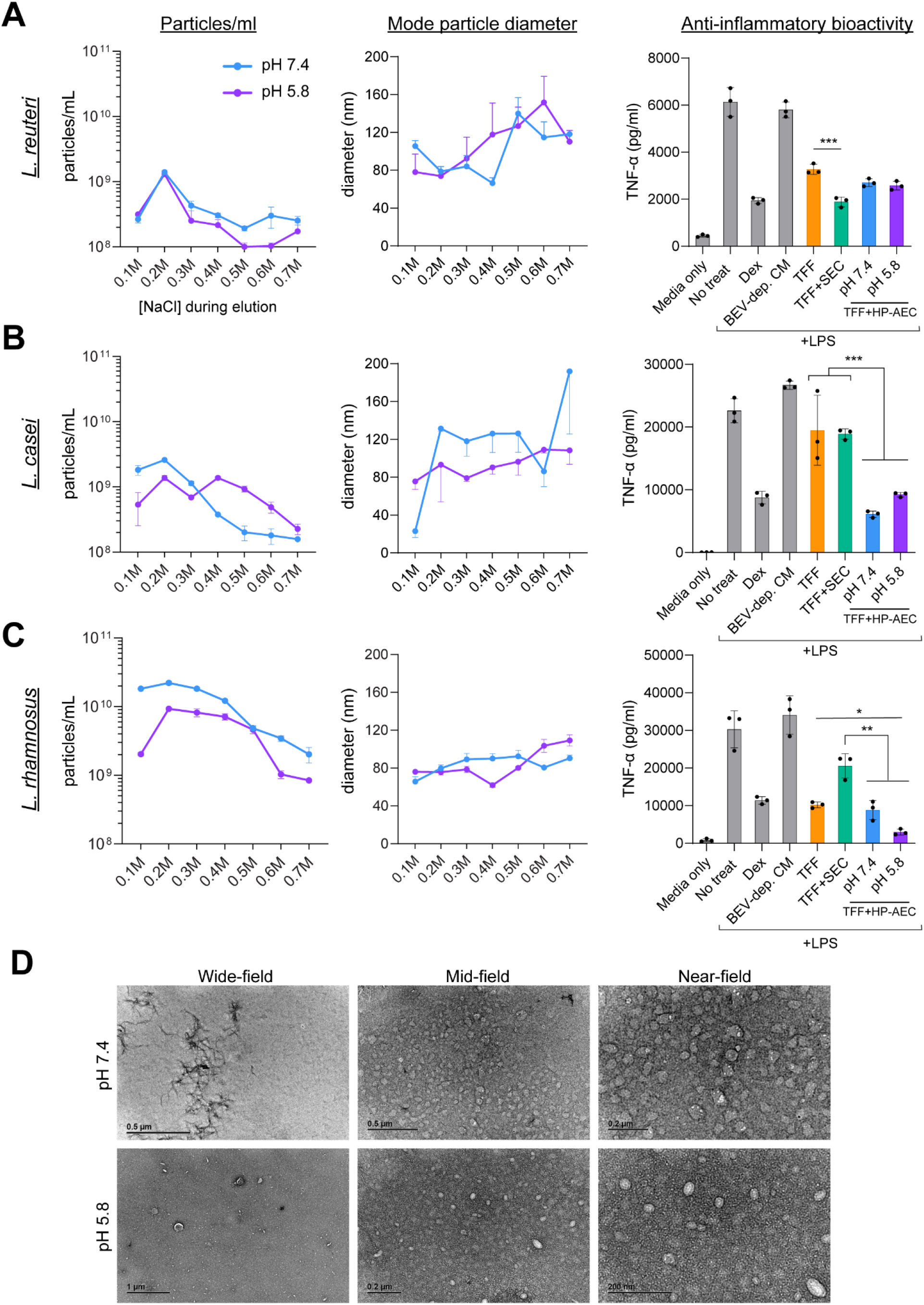
HPAEC can be applied for separation of BEVs from Gram-positive lactic acid probiotic bacteria. **A-C)** Nanoparticle tracking analysis (left) and anti-inflammatory bioactivity in an LPS-stimulated RAW264.7 mouse macrophage assay (right) of *L. reuteri* (A), *L. casei* (B), *L. rhamnosus* (C) BEVs following TFF + HPAEC purification at pH 7.4 or pH 5.8 (n=3 technical replicates). For the anti-inflammatory assay, BEVs for all purification methods were suspended in 300 uL media at the following concentrations: *L. reuteri* 0.1 ug/mL; *L. rhamnosus* 0.1 ug/mL; *L. casei* 0.01 ug/mL (n=3). D) Transmission electron micrographs *L. rhamnosus* BEVs purified by TFF + HPAEC at pH 7.4 and pH 5.8 with wide-field (left), mid-field (middle) and near-field views (right). Statistical significance was determined by ordinary one-way ANOVA with Holm-Šidák post hoc test. * p<0.05, ** p<0.01 ***p<0.001, ****p<0.0001.

Most importantly for biomanufacturing of probiotic BEVs, we assessed anti-inflammatory effects in TFF + HPAEC purified BEVs via suppression of TNF-α secretion from LPS-stimulated mouse macrophages. BEVs from all three probiotics that were processed by any of the three methods (TFF only, TFF + SEC, and TFF + HPAEC) were anti-inflammatory, as expected, at low protein doses (0.01 – 1 ug/mL) (Figure 4A-C). In contrast, BEV-depleted conditioned media (TFF permeate) had no anti-inflammatory effects (Figure 4A-C), indicating anti-inflammatory bioactivity is concentrated within BEVs at these relatively low protein doses. Notably, we observed more potent anti-inflammatory effects from TFF + HPAEC-purified BEVs compared to those obtained by TFF only or TFF + SEC. For *L. casei,* HPAEC performed at either pH yielded BEVs with more potent anti-inflammatory effects compared to both TFF-only and TFF + SEC. Similar results were observed for *L. rhamnosus*, except HPAEC at pH 5.8 only produced BEVs with more potent anti-inflammatory effects than TFF-only (Figure 4B). This result provides further evidence of the potential for pH reduction as a method to optimize HPAEC purification of BEVs. Of note, no potentiation of anti-inflammatory effects was observed for *L. reuteri,* which could possibly be attributed to their exceptionally low BEV yields (Figure 4A).

## 4. Discussion

Our results demonstrate that HPAEC can be applied for charge-based purification of probiotic BEVs. Furthermore, size-based TFF in tandem with charge-based HPAEC was shown to reduce co-isolated protein impurities more effectively in BEV samples compared to TFF alone or TFF + SEC. In general, BEV purification methods have relied on separation by size and/or density ^28^. However, BEVs are heterogenous in size (20-200 nm) and density (1.1 – 1.2 g/mL), and these ranges overlap with abundant bacterial culture products such as protein aggregates, pili, and flagella ^4^. Differential ultracentrifugation has been widely employed for BEV isolation ^28^ and relies on low-speed centrifugation to pellet larger-sized cells and debris and a subsequent high-speed centrifugation (typically >100,000g x 1-2 hours) to pellet BEVs. However, this method often co-isolates numerous contaminants and is generally not scalable, necessitating development of alternatives. A subsequent purification step using density gradient ultracentrifugation can further purify BEVs based on density, but requires prolonged ultracentrifugation (>18 hours), is technically challenging, and is again, not easily scalable. More recently, TFF and SEC have emerged as alternatives for EV and BEV purification. However, these methods rely on size-based separation and accordingly are prone to the co-isolation of similar sized cellular structures and protein aggregates, as we have demonstrated (Figure 1A-E). Thus, our method here simultaneously addresses two major challenges in BEV purification, i) lack of scalability, and ii) co-isolation of protein impurities.

Critically, reducing co-isolated protein impurities may enhance therapeutic efficacy of probiotic BEV therapeutics. Since many probiotic BEVs exert therapeutic activity via suppression of pathologic inflammatory responses, we utilized an *in vitro* model of macrophage inflammatory responses to assess potentiation of BEV bioactivity. Here, we confirmed prior reports that BEVs derived from lactic acid bacteria (*L. casei, L. reuteri, L. rhamnosus*) have anti-inflammatory effects in LPS-stimulated macrophages ^27,29^. Furthermore, at relatively low protein doses (less than 1 ug/mL BEV protein), we show anti-inflammatory bioactivity from probiotic conditioned media is associated with BEVs by using BEV-depleted conditioned media controls (Figure 4A-C). This allows anti-inflammatory bioactivity to serve as a surrogate identifier for Gram-positive BEVs, despite the lack of specific BEV markers. That is, higher ratios of BEVs to total sample protein will produce greater anti-inflammatory effects at sub-1 ug/mL doses in RAW264.7 macrophages. In accordance, our data support the conclusion that TFF + HPAEC also improves purity of Gram-positive probiotic BEVs due to potentiation of anti-inflammatory effects with TFF + HPAEC isolated BEVs compared to TFF or TFF + SEC (Figure 4A-C). As such, our method can potentially produce more potent probiotic BEV therapeutics via improving BEV purity – which is clearly desirable in biomanufacturing.

Flagellin presents purification challenges not only due to its similar size as BEVs when assembled into flagella, but also due to its nature as a negatively charged protein at physiologic pH. Indeed, we found that both flagellin and BEVs were bound to AEC resin (Figure 2G). We tested two solutions to this problem, i) the use of specific ionic strength buffers to elute flagellin and BEVs at different times, and ii) direct alteration of the binding affinity of flagellin and BEVs to the AEC column via pH reduction (Figure 3). Notably, both solutions resulted in improved EcN BEV purity. Furthermore, for EcN, the former approach yielded exceptionally pure BEVs and may be a promising approach for -omics studies. The latter approach of pH adjustment produced 2-fold increased yields of EcN BEVs with purity greater than TFF or TFF + SEC, and potentiation of *L. rhamnosus* BEV anti-inflammatory bioactivity (Figure 4), suggesting promise for biomanufacturing. Finally, similar approaches may be applicable to mammalian EV purification by AEC with respect to albumin, which is a common co-isolated impurity and is negatively charged.

The current lack of reliable probiotic BEV biomarkers complicates analysis of BEV purification. In this study, we relied on i) nanoparticle tracking analysis, which is not specific for BEVs and also detects protein aggregates, ii) qualitative electron microscopy, and iii) anti-inflammatory bioactivity assays which can be confounded by co-isolated anti-inflammatory molecules. Thus, biomanufacturing of Gram-positive BEVs would benefit from identification of novel critical quality attributes (CQAs), such as specific BEV protein markers, to support improved quality control processes. Additionally, clinical translation of BEVs will necessitate high yield BEV production, however, multiple purification steps as proposed here will reduce BEV yields. Since we and others have observed conventional chromatography resins have relatively poor BEV binding capacity compared to proteins ^30^, development of chromatography resins optimized for BEVs is needed. Along these lines, we did not optimize BEV loading quantity nor the reduced pH value of 5.8, which was selected based on literature values of mammalian EV isoelectric points and predicted isoelectric point of flagellin ^31^. Thus, optimization of HPAEC parameters, identification of CQAs for Gram-positive BEVs, and methods to improve BEV yields, especially for Gram-positive probiotic BEVs, will benefit BEV biomanufacturing.

## 5. Conclusions

In summary, we demonstrated that size-based purification alone co-isolates protein impurities, in accordance with prior reports. Then, we showed that AEC can be used to isolate BEVs derived from Gram negative E. coli and Gram-positive LAB probiotics, owing to their negative surface charge. Most importantly, we developed a solution for co-isolation of protein impurities through orthogonal size- and charge-based BEV purification with TFF + HPAEC. We found that our method can dramatically improve BEV purity with promise to potentiate probiotic BEV anti-inflammatory efficacy. Since our method uses scalable techniques already widely employed in industry, we believe this could be readily adopted for large-scale BEV biomanufacturing.

## Acknowledgments

This work was supported by the National Institutes of Health (HL159590), the National Science Foundation (CBET 1932963, MCB 2227598, CBET 1750542), the Food and Drug Administration (U01FD005946), and by the University of Maryland via an Mpower Graduate Fellowship to NHP.

